# Paternity sharing in insects with female competition for nuptial gifts

**DOI:** 10.1101/2022.03.26.485936

**Authors:** Jessica H. Browne, Darryl T. Gwynne

**Affiliations:** Department of Ecology and Evolutionary Biology, University of Toronto Mississauga, Mississauga, Ontario L5L 1C6, Canada

**Keywords:** parental investments, sperm competition, nuptial gifts, female competition, paternity

## Abstract

When males contribute costly investments to offspring (i.e., paternal care or nuptial gifts), they are expected to have high confidence of paternity. While studies of species with exclusive male parental care have provided support for this hypothesis, paternity may be more uncertain in systems where males feed each mate. While mating co-occurs with each oviposition in many male-care systems, allowing control over insemination, mating is separated from egg-laying in nuptial-gift systems so that males typically compete more with rivals for fertilizations. Given the costly gift investment, and long refractory periods that limit male ability to gain fitness from frequent copulations, there should be strong selection to avoid cuckoldry, especially complete paternity failure. Further, because females are expected to mate frequently to obtain male-supplied nutrition prior to oviposition, males may be unable to avoid sperm competition because virgins will be rare (and lower quality), and females are expected to undermine male traits that limit remating or manipulate sperm stores. Thus, paternity sharing, in which males are not excluded from siring offspring completely, is an expected outcome of sperm competition in these systems. Using wild-caught females in two species (an orthopteran and a dipteran), in which sexually selected females compete for important male nuptial gifts, we examined paternity patterns and compared them to findings in other insects. Using microsatellite analysis of offspring and sperm stores, we found evidence of shared paternity in both study species, where very few males failed to sire offspring. Although paternity was not equal among sires, our estimates of paternity bias were similar to other insects with valuable nuptial gifts and contrasted with patterns in gryllid crickets where males supply little more than sperm and are frequently excluded from siring offspring. Further support for paternity-sharing is the lack of last-male sperm precedence in either of our study species.

## Introduction

When males provide their mates with nutrition or offspring care, females usually mate multiply to obtain these benefits (Gwynne and Simmons 1990; Herridge et al. 2016; Bondurianksy 2001). These investments can both have high costs for males, and be essential for offspring production, leading females to compete for males and sometimes experience greater sexual selection relative to males (Tobias et al. 2012; Hare and Simmons 2018) (Trivers 1972; Thornhill 1979; Gwynne 1981; 1991; 2016; Gwynne and Simmons 1990; Simmons 1992; Herridge et al. 2016). In species with costly paternal care (many fish and birds; Clutton-Brock 1991), female re-mating rates are limited not only by the cost of eggs but also by the availability of parental males, which can drive fierce competition for the few mating opportunities. For example, female Gulf pipefish (*Syngnathus scovelli*) compete for 0-4 mates (mean: 1.6-2.2) and are subject to strong sexual selection on ornaments that attract males (Jones et al. 2001; Flanagan et al. 2014). Caring males typically have high confidence in paternity of the offspring they invest in, either due to control over insemination such as when unfertilized pipefish eggs are placed in an enclosed male brood pouch (Jones and Avise 2001) or when there are repeated copulations prior to each egg laid (*Abetus* water bugs; Smith 1979a, and sequentially polyandrous birds; Møller and Birkhead 1991; Schamal et al. 2004; Owens et al. 1995; Delehanty et al. 1998). This evidence has been central to parental investment theory (e.g., Trivers 1972; Westneat and Sherman 1992; Møller and Birkhead 1993; Requena and Alonzo 2017), which suggests that high paternity confidence facilitates the evolution of paternal care, given the fitness costs of cuckoldry (investment without paternity).

Paternity confidence seems to be more uncertain in systems where males provide nuptial gifts (common in insects: Thornhill 1976; Lewis et al. 2014) because there is less male control over insemination. As insect copulation typically occurs separately from egg-laying, mate (and thus paternity) monopolization by each of a female’s mates does not co-occur with each copulation/oviposition episode as in many male-care species. Thus, when female insects mate multiply, each ejaculate competes with the spermatheca-stored sperm of rivals for fertilizations, which can lead to biased paternity shares that is the essence of sperm competition theory (Parker 1970; Simmons 2001). Sperm competition risk and intensity might be especially high in species with mating gifts, as females have a strong motivation to mate frequently (Gwynne and Simmons 1990; Herridge et al. 2016) and may be less constrained by male availability relative to species with male care; while males providing nuptial gifts have refractory periods of a few days (e.g., the katydid *Requena verticalus*: Gwynne 1990), caring males typically have prolonged non-mating periods: male pregnancy lasts several-weeks in pipefish (Berglund and Rosenqvist 2003) and male *Abetus* water bugs carry eggs on their back for nearly a month until they hatch (Smith 1979b). Indeed, females have been observed mating with many males in several species with nuptial gifts, while virgin females are rare (e.g., 1-7 mates in *Rhamphomyia longicauda* dance flies; Herridge 2016, 0-6 mates in *Hemiandrus pallitarsis* weta (Browne 2021), and 1-6 in *Pholidoptera griseoaptera* katydids; Parker et al. 2017).

Paternity bias, which may include unequal paternity among sires (paternity skew) and/or paternity failure, in which some males are completely excluded from fertilizing offspring (e.g., only 51-66% of a female’s mates sire offspring in gryllid crickets: *Laupala cerasina*; Turnell and Shaw 2015), occurs when the ejaculates of certain males have an advantage over others. In insects, paternity bias is frequently determined by mating order such that the last mating male fertilizes more of a female’s offspring than his rivals (Parker 1970; Lloyd 1979, Eberhard 1985; Simmons 2001; Parker et al. 2017). This is due to processes such as (1) sperm stratification within the spermatheca, layering of the sperm that results in the most recent sperm being used first to fertilize eggs, (2) sperm displacement or removal where a male’s sperm replaces that of past mates, and (3) mating plugs or guarding that reduce the probability of female remating. Compared to species where males contribute only sperm, males that invest heavily in mate feeding will experience high costs in paternity losses (cuckoldry: Trivers 1972), especially if their ability to mate again is limited by multi-day refractory periods (Gwynne 1990; Perry and Tse 2013). Not only are the costs of cuckoldry high when males invest in costly nuptial gifts, but female reliance on male gifts for survival or egg development probably undermine the ability of males to inhibit female remating or manipulate her sperm stores. Furthermore, males supplying valuable gifts are probably less able to avoid or reduce sperm competition by seeking unmated females (see Simmons 2001; Thomas and Simmons 2007) given the scarcity, and potentially low fecundity of virgins when females mate frequently to obtain nutritious gifts (Simmons 2007; Herridge 2016, Browne 2021, Parker et al. 2017).

In contrast, if there is shared paternity (presumably via sperm mixing and/or female control; Simmons 2001), each investing male will only sire a portion of a female’s brood, and there is a reduced chance of complete paternity loss. Given that the costs of siring no offspring are expected to be high in paternally investing systems, shared or near-equal paternity may benefit male fitness by increasing the probability of siring some of a female’s lifetime offspring production. While this is contrary to predicted increased paternity bias through last male sperm precedence in species with larger nuptial gifts (Gwynne 1984a; Simmons 2001), this prediction of shared paternity is not unlike paternity patterns in many parental care systems. Although caring males typically have high confidence of paternity in the offspring they care for, this does not represent a female’s lifetime production of offspring; subsequent eggs she lays are fertilized by other caring males. Indeed, female pipefish (*Syngnathus typhle*) distribute their eggs between several males, often within the same laying bout (Berglund et al. 1988).

Although studies of paternity in wild arthropods are limited, microsatellite analysis of offspring has revealed evidence of shared paternity in two nuptial gift-giving insects: a katydid, *Requena verticalis*, and a ladybird beetle, *Adalia bipunctata*. In both species, paternity was shared (multiple paternity) in nearly all broods, and importantly, it appeared that all males that mated with a female sired offspring (Simmons 2007; Haddrill et al. 2008). However, despite the low rates of paternity failure, paternity was still skewed in *R. verticalis* (not measured in *A. bipunctata)*, as males varied in the number of offspring sired (Simmons 2007). Similarly, in *Ephippiger ephippiger*, a katydid species where females can compete for spermatophylax gifts (Ritchie et al. 1998), each comprising about 20% of a male’s weight (Vahed and Gilbert 1996), paternity was distributed unequally among sires (Hockham et al. 2004). Although the authors concluded there is a high degree of paternity bias in this species, there was no estimate of female mate number, meaning they could not determine whether any males were excluded from fathering offspring.

A high degree of paternity skew and paternity failure have been reported in several wild-caught species of gryllid crickets (Simmons and Beveridge 2010; Turnell and Shaw 2015) that lack costly nuptial gifts, further supporting the idea that shared paternity may be an adaptive outcome of sperm competition when females compete for male-supplied nutrition. On the other hand, a similar level of paternity bias was found in *Pholidoptera griseoaptera*, a katydid species where males provide small yet substantial spermatophylax gifts (about 7% of their weight; Vahed and Gilbert 1996). Using the number of spermatodose sperm sacs to estimate female mating frequency, Parker et al. (2017) found unequal paternity shares among sires, and also a large proportion of males that sired no offspring. However, paternity estimates in this study were potentially limited. As *Pholidioptera* (as well as *Ephippiger*) require several winters for eggs laid late in the season to hatch (Hartley and Warne 1972; Ingrisch 1986), paternity analyses were conducted only on a portion of the offspring that reached the whole-embryo stage in the lab. Eggs requiring a diapause to continue development (about 60% of viable eggs) would not have been included (Parker et al. 2017) and this may have inflated the number of paternity-losing males (Gerlach et al. 2012; Fritzche and Arnqvist 2013).

Our study measures the proportion of males failing to sire offspring and the skew among sires in an orthopteran where, for the first time, we assess paternity in all of a female’s eggs, her lifetime reproduction. We also test for evidence of paternity sharing in a well-studied dance fly species in which ornament-displaying females compete for costly male gifts of prey that are necessary for offspring development.

In the New Zealand “short-tailed” ground weta, *Hemiandrus pallitarsis* (Orthoptera: Anostostomatidae), females mate frequently and obtain spermatophylax meals (adhered to her mid-abdomen, separate from the sperm-containing ampulla; Gwynne 2002; 2004), probably to allow them survive a period of 5-6 months without food while caring for offspring before dying (Gwynne 2004). At the end of a mating season females lay all their eggs in a brood chamber (hence their “short-tail” reduced ovipositor) potentially reducing a male’s chances of gaining paternity via last male sperm precedence (most insects have multiple ovipositions). In *H. pallitarsis*, there is evidence that females are under sexual selection, which may explain an “ornamental” secondary genitalic device (accessory organ: Gwynne 2004, 2005), that is inserted between two main parts of the male genitalia as the gift is deposited (Gwynne, unpublished). Sexual selection on female accessory organs may be a result of males mating with longer accessory organs (Browne 2021), possibly because it signals higher fecundity (Gwynne 2005). Alternatively, longer organs may increase copulation duration thus allowing larger nuptial gifts to be obtained, as suggested for the function of the “female penis” in certain psocopterans (Yoshizawa et al. 2014; Yoshizawa et al. 2018) and the ridges on the subgenital plates of some katydids with sexual competition among females (Gwynne 2001).

In the empid dance fly (Diptera: Empidae), *Rhamphomyia longicauda*, males provide females with prey items (usually small insects) in exchange for mating. Females rely on these mating gifts to obtain protein for egg development (Downes 1970; Hunter and Bussière 2019) and mate multiply, with an estimated 1-7 males (mean: 2.5; Herridge 2016). Females possess two sex-specific ornaments (pinnate scales on the legs and inflatable abdominal sacs) that function in attracting prey-carrying males in mating swarms at dusk or dawn. Although studies have not shown directional sexual selection on females in this species (See Wheeler et al. 2012; Herridge 2016), there is evidence that males prefer females with larger ornaments (Funk and Tallamy 2000; Murray et al. 2018), even though ornament size correlates weakly with egg number and size (Wheeler 2008; Funk and Tallamy 2000). While we were unable to determine paternity for a female’s lifetime reproduction, *R. longicauda* is ideal for testing whether paternity is shared, since females possess a single sclerotized spermatheca, creating high potential for sperm displacement and thus biased paternity in favor of the last male (Simmons 2001). Further, because females rely on prey gifts to complete ovarian development (Downes 1970; Hunter and Bussière 2019), male behaviours that reduce the probability of future matings (gift acquisition) by females are unlikely to be adaptive for either sex, due to the fitness of their offspring being compromised. In both these species, we expect lower paternity bias and less last-male sperm precedence compared to most insects. We predict similar paternity shares among sires, and importantly, that few males will fail to fertilize offspring due to the fitness costs, given their valuable investment.

## Materials and Methods

### Collection and rearing

We collected 17 *H. pallitarsis* pairs (females and their (last) mate guarding male) in 2017 from Kiriwhakapapa trail near Masterton, NZ (-40.807627, 175.546532) and two private gardens in Palmerston North, NZ (-40.413909, 175.662814). Another 10 pairs were collected in 2002 by D. Gwynne from the same locations. Males were preserved at -20°C for DNA analysis and live females were transported to our lab in Canada to oviposit eggs. This was an extended process that involved placing females in artificial brood chambers formed in potter’s clay (2002) or soil (2017) and allowing 2-3 months for females to lay eggs, as well as an additional 5-6 months for eggs to mature. While in the brood chambers, we exposed females to typical winter soil temperatures, which were based on New Zealand weather records for March to September. All but 3 females laid eggs, however these only developed (hatched nymphs or visible eye spots) in 19 of the broods. After eggs began hatching, we collected the offspring from 17 broods (excluding two where less than 5 offspring developed) and preserved them along with their mothers at -20°C. We dissected spermathecae from females and isolated the sperm contents using >70% ethanol, which causes sperm to harden into a pellet (Tripet et al. 2001) and stored them at -20°C.

Similarly, we collected *R. longicauda* mating pairs (in copula, n=131) from mating swarms near the Credit River, Glen Williams, Ontario, Canada (43.6865660, -79.9260960) from mid-June to mid-July of 2017. Males were killed and preserved via freezing while females were placed in individual plastic containers to lay eggs. They were provided with a variety of potential oviposition substrates as oviposition location in the wild is unknown. Mortality was high (88 females died prior to oviposition), however, 43 females laid 1-2 batches of eggs (1-89 eggs overall) on the sides of the containers before eventually dying. Dance fly eggs, which do not require maternal care for survival, were collected and stored in petri dishes lined with damp filter paper. They were left to develop at room temperature for up to six days, at which point those showing signs of maturation (i.e., darkening of the chorion, visible mouthparts, hatching) were frozen for genetic analysis. We removed female’s spermathecae and again used strong ethanol (>70%) to isolate the sperm pellet from female tissue (as in Tripet et al. 2001).

### DNA Extraction and Microsatellite Analysis

DNA was extracted from *H. pallitarsis* adult and offspring tissue using a Proteinase K based extraction method with multiple ethanol washes. DNA was extracted from sperm pellets using a Qiagen DNeasy Blood and Tissue Extraction kit. We followed a modified version of the recommended protocol, which involved adding 12µL DTT to each sample to improve lysis of sperm cells and dissolving DNA in a reduced volume (50µL) of elution buffer. Unfortunately, DNA extractions proved to be much more difficult in dance flies. While we were able to utilize the same Proteinase K based method to extract DNA from the bodies of *R. longicauda* females and their most recent mate, we had difficulty extracting sufficient quantities of DNA from offspring (developed eggs or first instar larvae). We eventually had some success using a Chelex-based extraction with an initial homogenization step; however, we were unable to extract DNA from sperm pellets in quantities sufficient for PCR, despite using a QIAamp DNA Mirco kit with the addition of DTT as in Herridge (2016).

In both ground weta and dance flies, we genotyped samples at four loci using fluorescent labelled microsatellite markers. Markers for *H. pallitarsis* were developed by Genetic Marker Services, Brighton, UK (table 1) and characterized in two populations (see Browne 2021). We conducted PCR in two multiplex reactions, each containing 5.75µL sterile filtered water, 1µL 10X PCR reaction buffer, 0.2µL 10mM dNTP mixture, 0.5µL of forward primer (with fluorescent labels HEX or 6-FAM), 0.5µL reverse primer, and 1µL of template DNA. Thermocycling was done using an Eppendorf Mastercycler with the following temperature regime: 2 minutes at 94ºC followed by 35 cycles of 30 seconds at 94ºC, 30 seconds at 60ºC, and 15 seconds at 72ºC, then an additional 3 minutes at 72ºC.

**Table 1.**
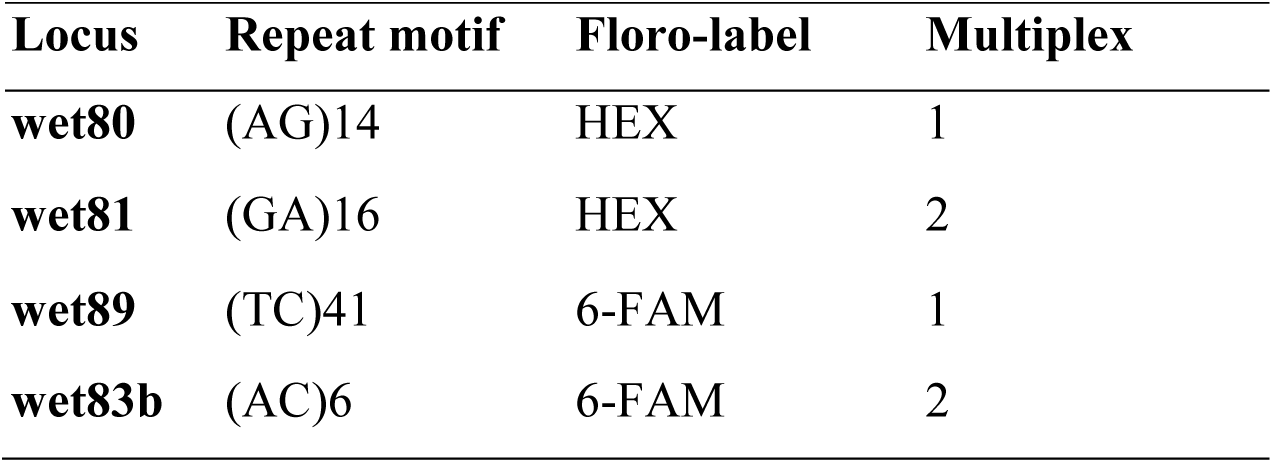
Microsatellite loci used to genotype *H. pallitarsis* (See Browne 2021 for details)

Markers for *R. longicauda* were developed and characterized by E. J. Herridge and L. Bussière at University of Stirling for use with the same study population (table 2; Herridge 2016). We again conducted PCR in two multiplex reactions, this time each containing 5µL sterile filtered water, 1µL 10X PCR reaction buffer, 0.1µL 50M MgCl_2_, 0.2µL 10mM dNTP mixture, 0.75µL forward primer (with fluorescent label on 5’ end), 0.75µL reverse primer, and 2µL of template DNA (1µL for adults). The thermocycling regime was adapted from Herridge (2016) and included 5 minutes at 95ºC followed by 40 cycles of 94ºC for 30 seconds, 56 ºC for 30 seconds, 72ºC for 25 seconds, then an additional 72ºC for 10 min. We sent all PCR products to The Centre for Applied Genomics in Toronto, Ontario for fragment analysis on an Applied Biosystems 3730xl capillary sequencer. The resulting electropherograms were examined using GeneMarker V1.97, which allowed us to identify fluorescence peaks, representing alleles.

**Table 2.**
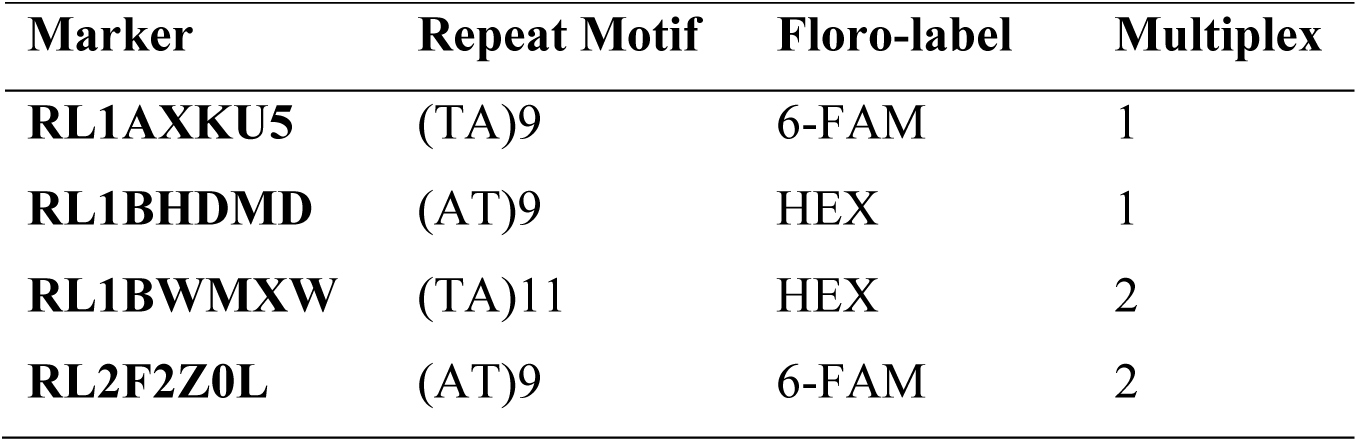
Microsatellite markers used to analyze *R. longicauda* (See Herridge 2016)

### Paternity Analysis

Because of difficulties genotyping dance fly offspring, methods of paternity analysis differed between the two species. In ground weta, we used data from 11-51 (mean: 28.8) developed offspring (95% of those collected) to estimate the number of sires in each brood. We first estimated the minimum sire number using GERUD 2.0 (Jones 2005), a parentage program that computes the most likely minimum father combination given the offspring genotypes across multiple loci. Offspring that shared an allele with a female’s last mate at all loci were considered to be fathered by this male. Additionally, we estimated the most likely number of sires in each brood using Colony Version 2.0.6.6 (Jones and Wang 2010). This program uses population allele frequencies (determined in Browne 2021) to determine the most likely parental configuration. A female’s last mate was included in Colony as a candidate father to determine which offspring were most likely sired by him (See Turnell and Shaw 2015 for comparison of parentage programs).

We then estimated the number of males that failed to sire offspring by identifying alleles in a female’s sperm storage organ that were not present in any of the offspring. We used allele counting (assumes heterozygosity; number of unique paternal alleles present in offspring divided by 2) to estimate the minimum number of additional males for each brood. In the dance flies, our analysis was more limited. While we genotyped 11-58 (mean: 32.6) of a female’s lab-laid eggs, this only accounted for a portion (45% on average) of those collected, and in some cases only included two loci. Further, we could not determine which male alleles remained in the spermatheca, as we were unable to extract DNA from female sperm pellets. Using the available offspring data for 11 dance fly broods, we first counted the unique paternal alleles present in offspring and divided by 2 to estimate the minimum number of sires at each locus. Next, we determined the most likely sire configuration using Colony (Jones and Wang 2010), again including the female’s last mate as a candidate father. In the absence of DNA from female sperm stores, we compared our estimates of sire number to estimates of mate number for this population (determined by Herridge 2016) to assess whether males likely experience paternity loss. We used the estimate of minimum sire number (allele counting form offspring) so that this would be comparable to Herridge’s (2016) estimate of minimum mate number (allele counting from spermathecal stores).

### Statistical Analysis

In both ground weta and dance flies, we tested whether paternity patterns deviated from a “fair raffle” scenario (Parker et al. 1990), in which males sire equal (or near equal; Herridge 2016) proportions of offspring. We measured the evenness of paternity shares in each brood where females mated multiply by calculating the paternity skew and plotting this against sire number. For this we used Starr’s (1984) measure of paternity skew (Σ(proportion offspring sired)^2^ in which a value of 1 represents complete paternity bias in favor of a particular male. For both weta and dance flies, we determined whether the observed paternity skew was significantly higher than the skew expected if offspring were distributed evenly to all sires (as in Simmons and Beveridge 2010; Turnell and Shaw 2015) using the 95% confidence intervals of the best fit line as well as a paired t-test. Finally, we tested for evidence of last male sperm precedence in both *H. pallitarsis* and *R. longicauda* by determining whether a female’s last mate fathered a greater proportion of offspring than other males, or than would be expected by equal shares.

## Results

### Weta: Sire Number

Among the 17 *H. pallitarsis* broods, the minimum sire number (GERUD) averaged 3 ± 1.6 SD (range 1-6). Estimates of most likely sire number (Colony) were much higher with an average of 5.9 ± 2.4 SD (range 2-10). Regardless of the method used, the number of sires detected did not change with the number of offspring tested (GERUD: ß=0.03, R^2^=0.03, p=0.525, Colony: ß=0.08, R^2^=0.12, p=0.173). The majority of females mated multiply and had offspring sired by at least 2 males (GERUD: 83% Colony: 100%)

### Weta: Paternity Bias

Additional alleles, suggestive of males that mated but did not sire offspring, were found in the spermathecae of 5 females, however, these rarely represented more than 1 male per brood. On average, 85.4% (binomial 95% CI: 74.6%, 93.1%) of the males that inseminated a female fathered offspring. The rate of paternity success was similar (mean: 88.6%, binomial 95% CI: 81.9%, 93.5%) when using Colony to estimate sire number. Despite this, paternity shares among weta sires were still biased. Overall, the paternity skew (degree to which paternity is unevenly distributed) was significantly higher than would be expected if all sires fathered an equal number of offspring (one-tailed paired t-test, mean difference = 0.10, t = 3.72, df = 12, *p* = 0.001; fig. 3). This result was consistent when using Colony to estimate the most likely sire number (one-tailed paired t-test, mean difference = 0.09, t = 4.17, df = 16, *p* = 0.0004). Paternity skew declined linearly with the number of sires (GERUD: ß= -0.10, R^2^ = 0.61, *p* < 0.001; Colony: B= -0.03 R^2^ = 0.42, *p* =0.003), although this was not a better fit than an inverse (non-linear, 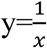) relationship (GERUD: R^2^ = 0.64, *p* < 0.001, *δ*AIC < 1.0; Colony: R^2^ = 0.43, *p* =0.002, *δ*AIC < 1.0).

**Figure 1.**
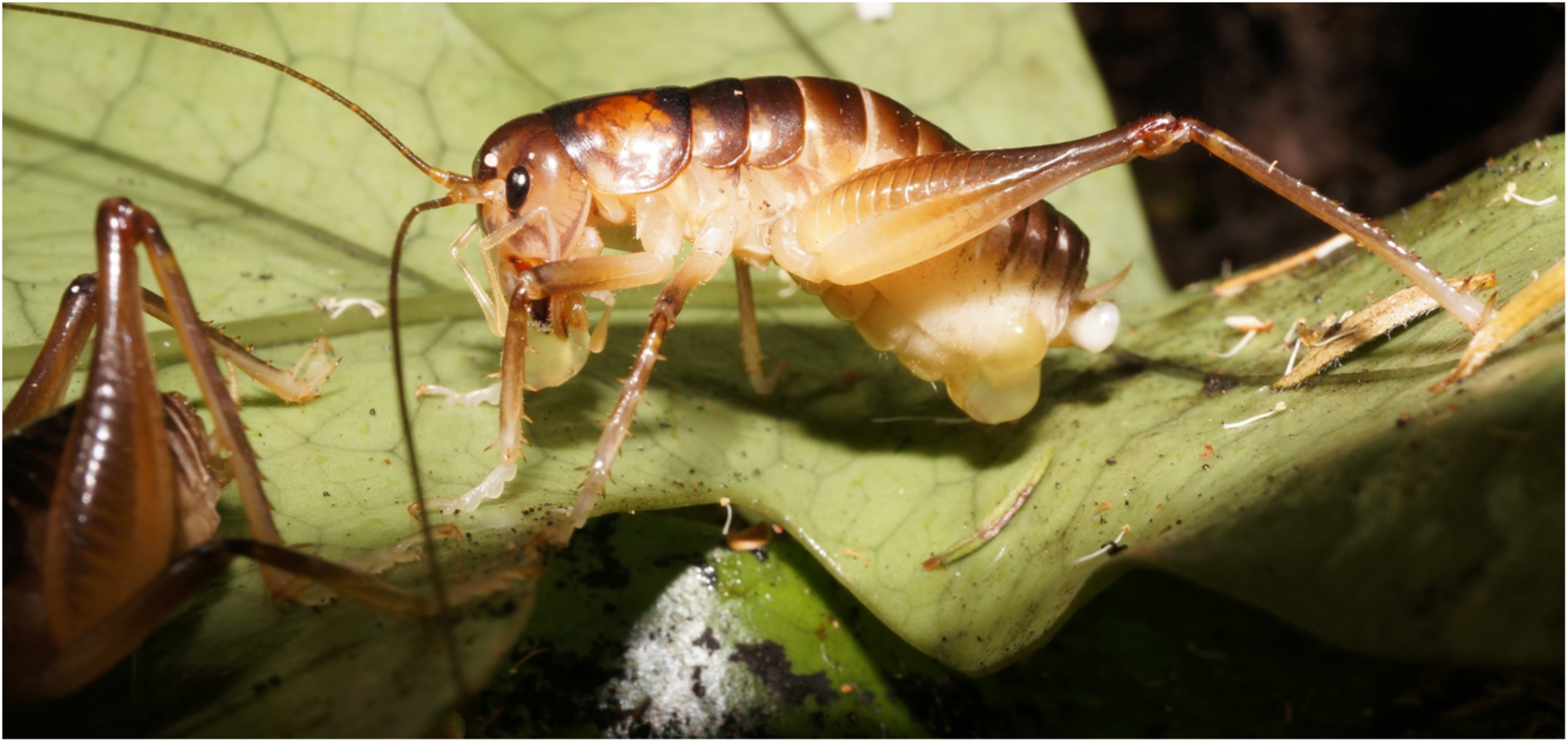
Female *H. pallitarsis* just after mating, eating part of the spermatophylax gift. One lobe remains adhered to her abdomen and the globular sperm package is visible at the end of her abdomen. Photo by Darryl Gwynne.

**Figure 2.**
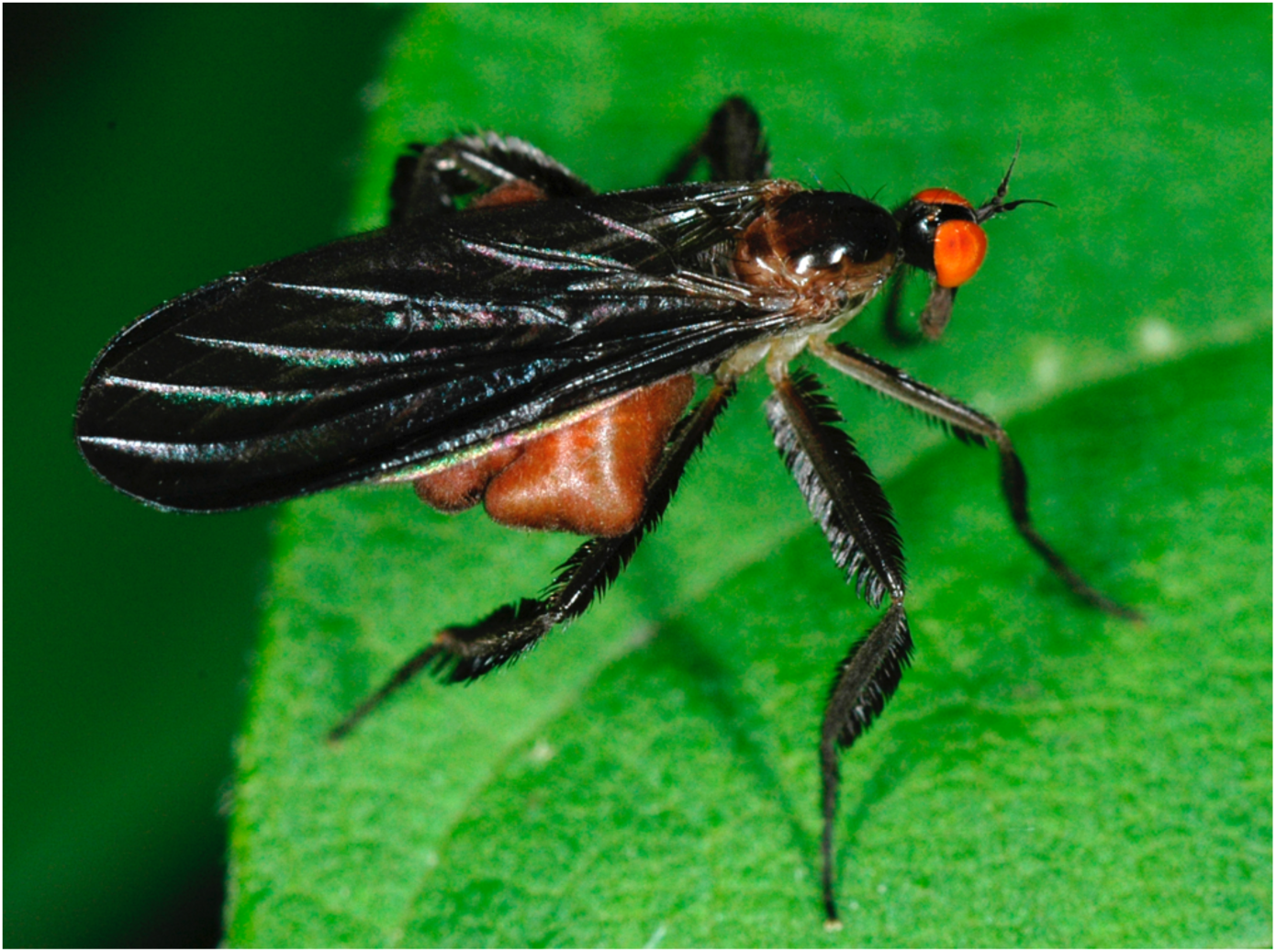
Female dance fly (*Rhamphomyia longicauda*) showing inflated abdomen and leg-scale ornaments. Photo by Heather Proctor.

**Figure 3.**
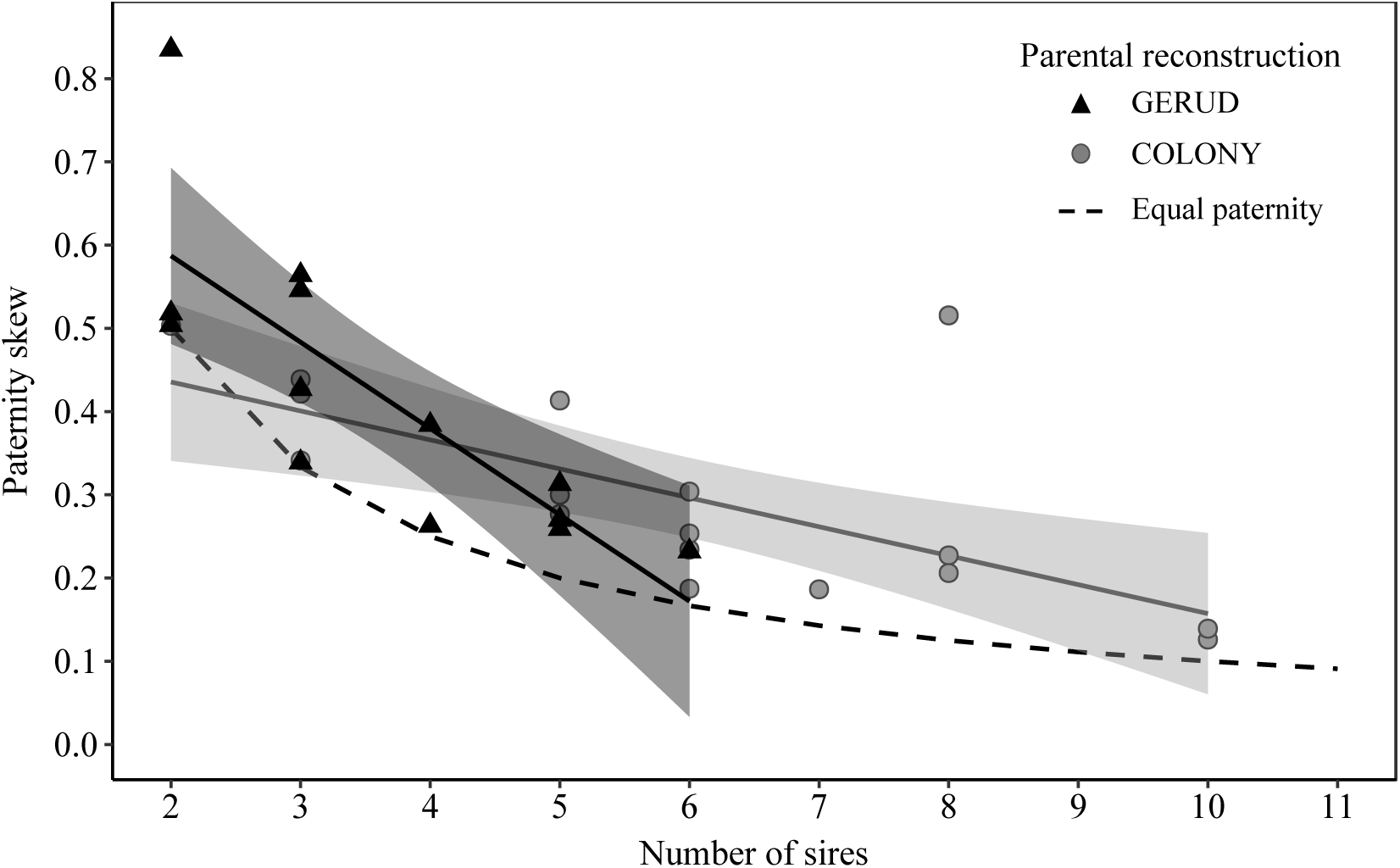
Paternity skew occurring in broods of multiply mated *H. pallitarsis* females, plotted against the minimum number of sires (parental reconstruction determined using GERUD; n=13) or most likely number of sires (Colony; n=17) estimated from microsatellite analysis of offspring. The regression lines (solid) show the relationship between number of sires and observed paternity skew with 95% confidence intervals showing deviation from the null skew expected when all sires father an equal number of offspring (dashed line).

### Weta: Last male sperm precedence

A female’s last mate did not consistently have a fertilization advantage. Based on the minimum father combination (GERUD), last males were seldom the most successful male (42% of broods) but fathered offspring in all but two broods (Appendix). Overall, a female’s last mate did not father a significantly greater proportion of offspring than previous males (Welch two sample t-test: t = 0.37, df = 16.89, *p* = 0.357) or that expected under a fair raffle scenario (equal shares among sires) (one-tailed paired t-test: t = -0.70, df =11, *p* = 0.751; fig. 4).

**Figure 4.**
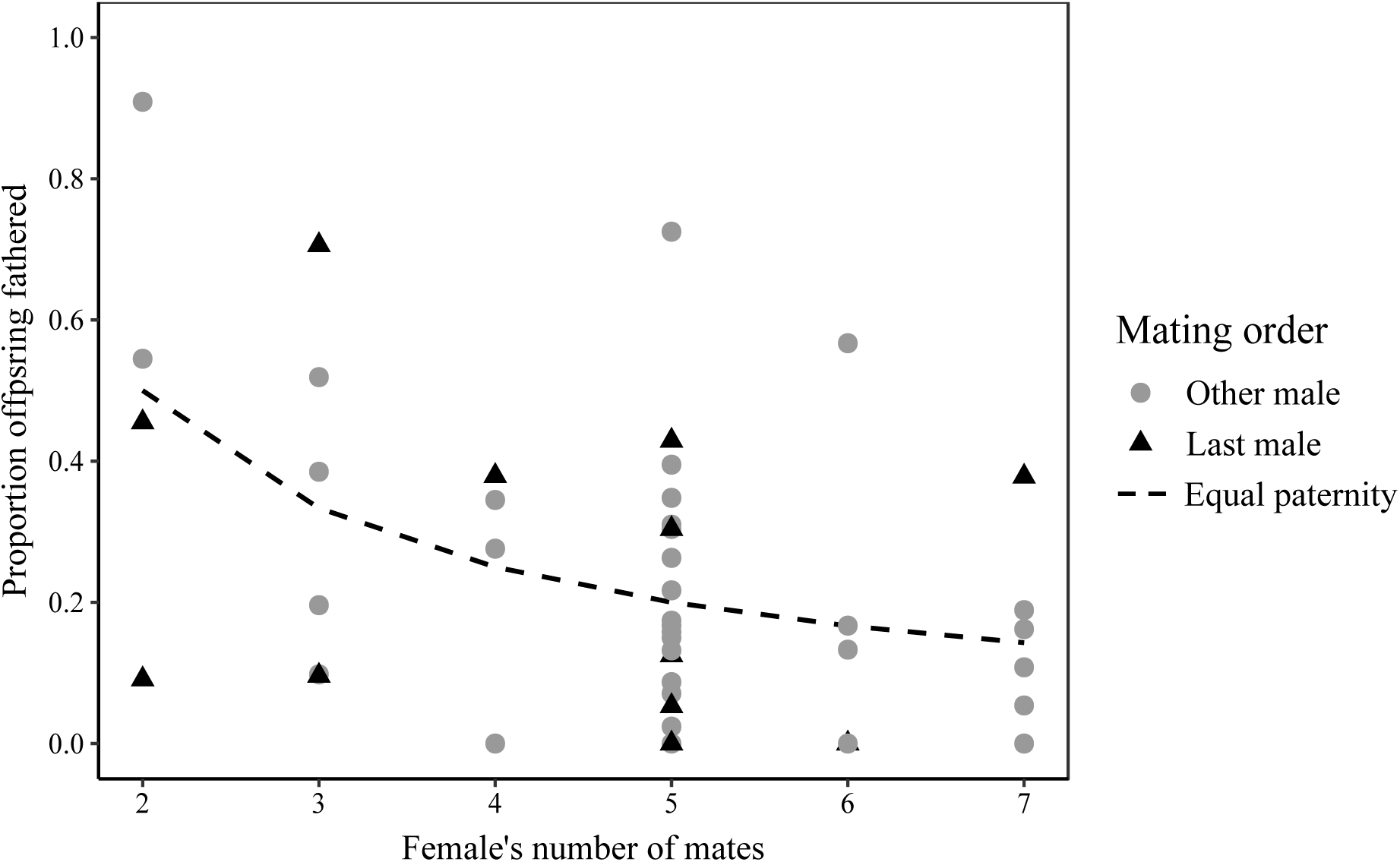
Proportion of offspring fathered by last male relative to all other competing males in *H. pallitarsis* ground weta (using GERUD – minimum sire configuration). Data are shown relative to the proportion of offspring each male would sire if paternity shares were equal (null paternity).

When using Colony to reconstruct the mostly likely paternal configuration, the last male was excluded from a further six broods (fathering the majority of offspring in only 24% of broods; Appendix) and overall did not father a significantly greater proportion of offspring than previous mates (Welch two sample t-test: t = 0.55, df = 17.75, *p* = 0.293) or that expected under a fair raffle scenario (equal shares among sires) (one-tailed paired t-test: t = -0.40, df =16, *p* = 0.651; fig. 5).

**Figure 5.**
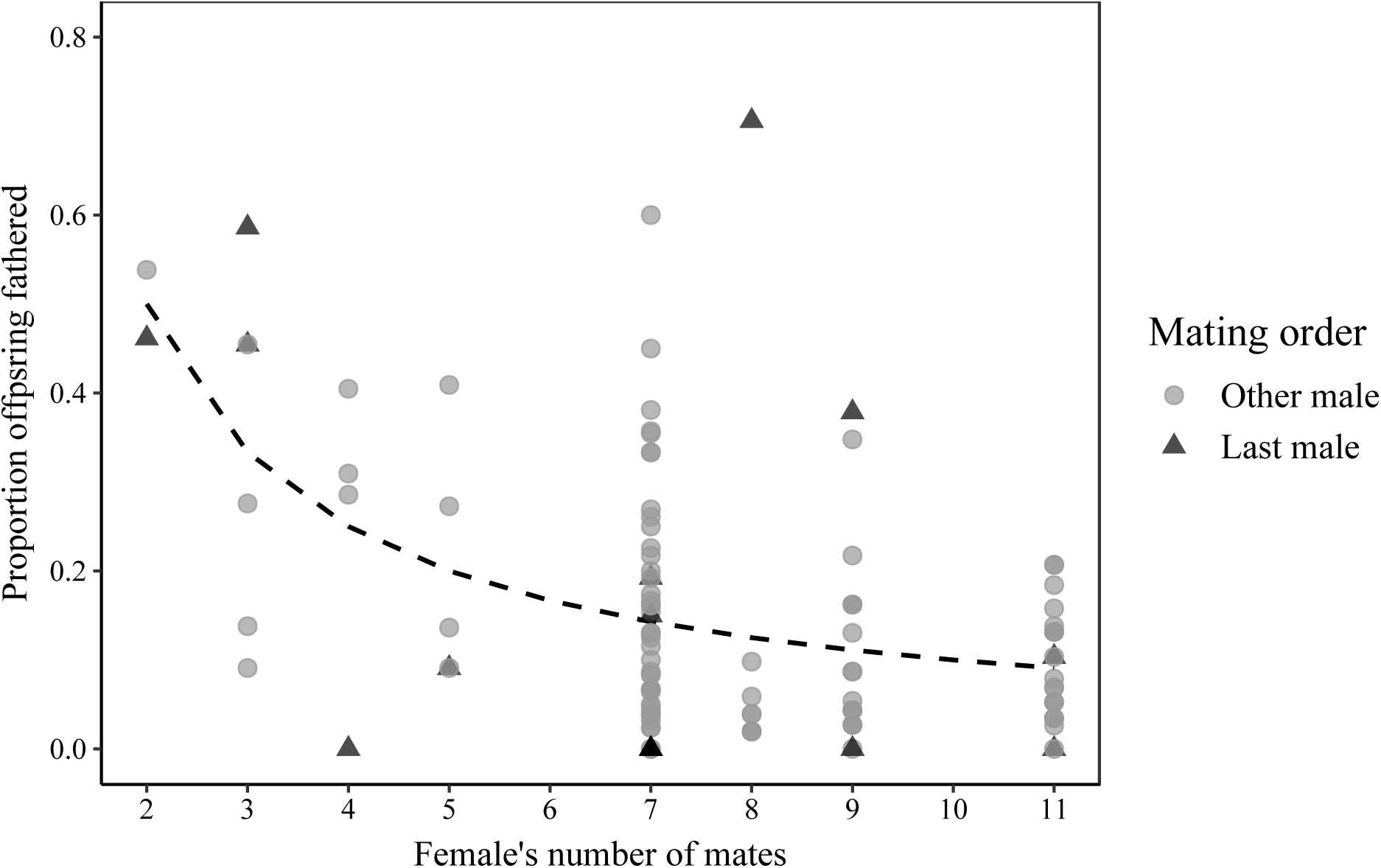
Proportion of offspring fathered by last male relative to all other competing males in *H. pallitarsis* ground weta (using COLONY – most likely sire configuration). Data are shown relative to the proportion of offspring each male would sire if paternity shares were equal (the null), indicated by the dashed line.

### Dance flies: Sire Number

We estimated sire number in dance flies using the 11 broods from which we were able to genotype offspring. When we used the conservative method of allele counting, the number of sires averaged 3.6 ± 1.3 SD and ranged from 2-6 sires. Estimates of sire number were much higher when using Colony, which averaged 13.5 ± 5.9 SD sires and ranged from 7-26. In dance flies, the number of offspring tested did influence the number of sires we were able to detect. We found a significant positive relationship between the number of offspring genotyped and number of sires detected for both of our estimates, however, this relationship was much stronger when using Colony rather than allele counting (Allele counting: ß=0.07, R^2^=0.69, p=0.002; Colony: ß=0.34, R^2^=0.87, p= <0.0001). Regardless of the method used to estimate sire number, all females mated multiply, and had offspring fertilized by at least two sires.

### Dance flies: Paternity Bias

Although we could not genotype the sperm stored in female’s spermathecae to estimate the proportion of failed matings, we found evidence of paternity skew among sires, similar to the weta. Using the most likely paternal configuration (Colony), the paternity skew was determined to be significantly higher than would be expected if all sires fathered an equal number of offspring (one-tailed paired t-test, mean difference = 0.03, t = 6.09, df = 10, *p* = <0.0001; fig. 6). Again, paternity skew declined linearly with number of sires (fig. 6; B= -0.004, R^2^ = 0.59, *p* < 0.003), although this was not a better fit than an inverse (non-linear, 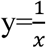) relationship (R^2^ = 0.57, *p* = 0.004, *δ*AIC < 2.0).

**Figure 6.**
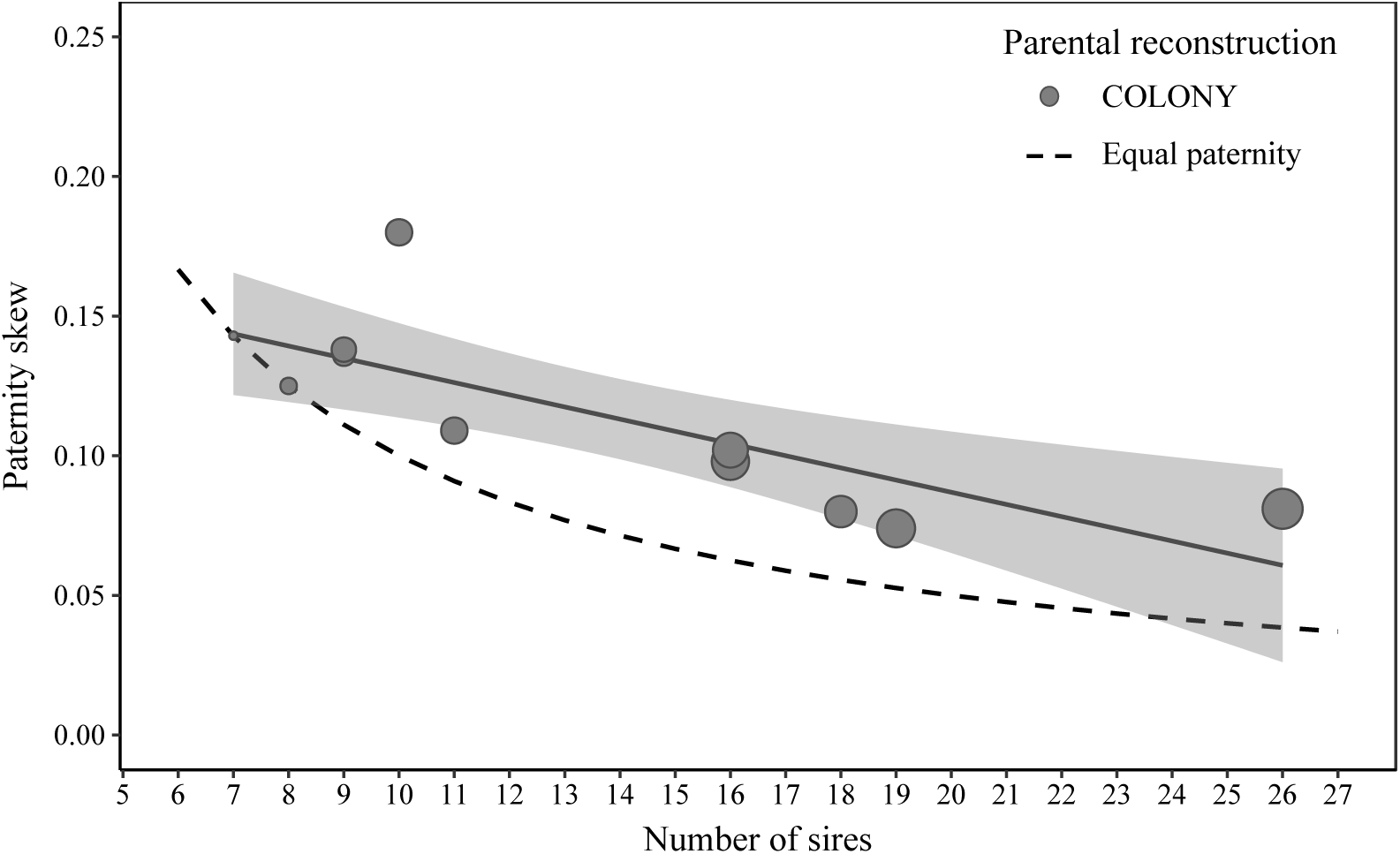
Paternity skew among offspring of female *R. longicauda* (n=11) plotted against the most likely number of sires (determined from microsatellite analysis of offspring; Colony). The regression line (solid) shows the linear relationship between number of sires and observed paternity skew with 95% confidence intervals showing deviation from the null skew expected when all sires father an equal number of offspring (dashed line). Size of points represents the number of offspring analyzed for each estimate of paternity skew, which has a positive effect on the number of sires detected (see main text).

### Dance flies: Last male sperm precedence

There was no evidence of last male sperm precedence in our samples. In some cases, the last male did not fertilize any offspring (Appendix), however it is unclear whether this is a result of sperm competition or our sampling methods, which may have disrupted mating pairs prior to full insemination. Overall last males fathered a significantly lower proportion of offspring than previously mated males (Welch two sample t-test: t = -4.07, df = 13.58, *p* = 0.001) and that expected under a fair raffle scenario (equal shares among sires) (one-tailed paired t-test: t = - 3.74, df =10, *p* = 0.004; fig. 7).

**Figure 7.**
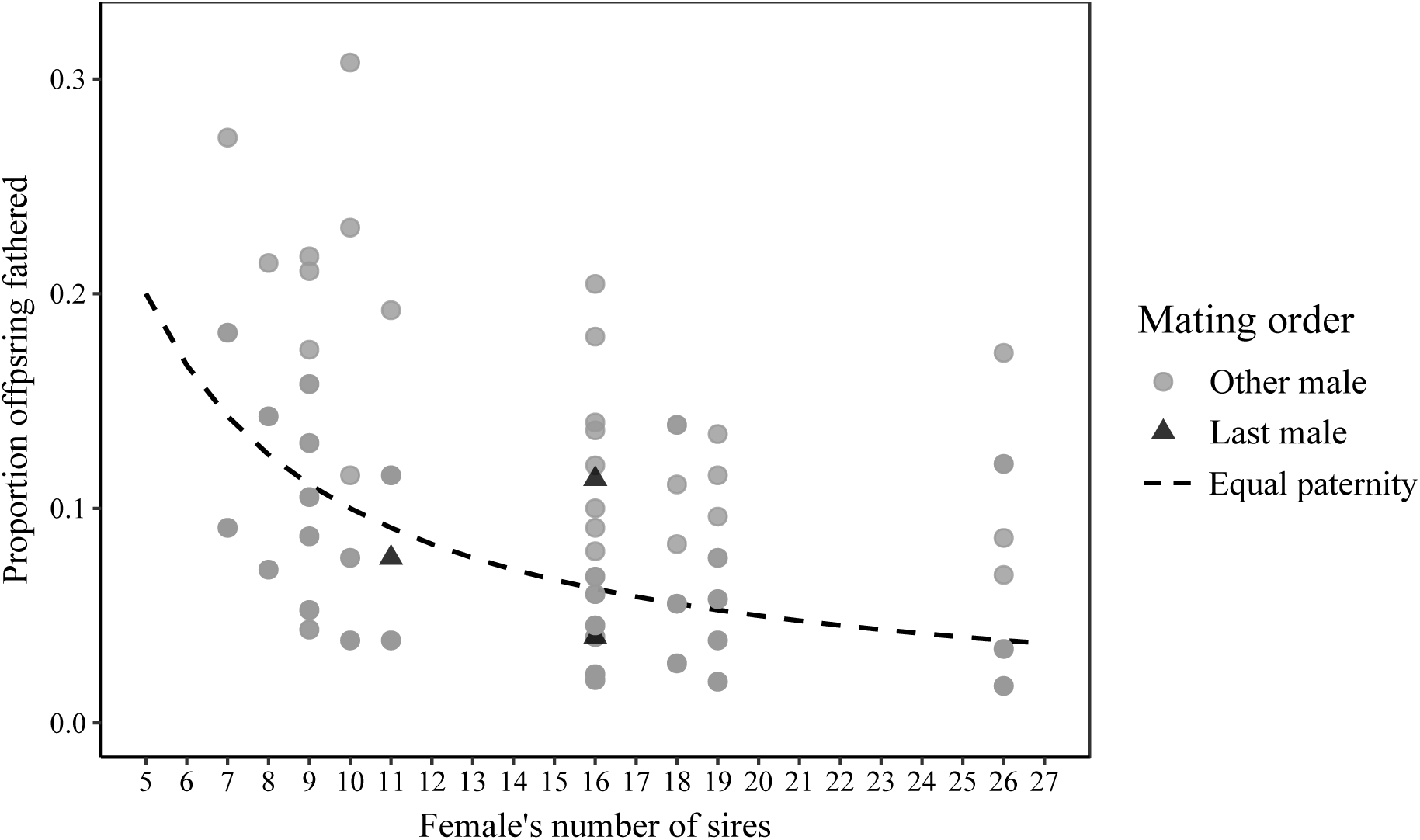
Proportion of offspring fathered by last male relative to all other competing males in *R. longicauda* dance flies (using COLONY – most likely sire configuration) Data are shown relative to the proportion of offspring each male would sire if paternity shares were equal (the null), indicated by the dashed line.

## Discussion

As predicted we show evidence of shared paternity (i.e., reduced paternity bias relative to other orthopteran insects; Simmons and Beveridge 2010; Turnell and Shaw 2015) in two insect species where ornamented females appear to compete for important nutrition provided by males during mating. In one of the species, we used female lifetime reproduction. While both species showed some degree of paternity skew among sires, paternity was shared among multiple males in most broods, and there were few males excluded from fathering offspring.

In ground weta (*H. pallitarsis*), paternity estimates using lifetime offspring production revealed a level of paternity confidence in which most investing males sired offspring. We found multiple paternity in most broods, with an average of 3.0 ± 1.6 SD (most likely estimate: 5.9 ± 2.4 SD) sires, and importantly, nearly all of the males that inseminated a female fathered offspring.

In dance flies (*R. longicauda*), multiple paternity was observed in all clutches, with offspring being shared between a high number of sires (minimum mean: 3.6, most likely mean:13.5). Although we could not measure the proportion of males that failed to sire offspring, it was assessed indirectly by determining if the number of mates (estimated in Herridge 2016 using same study population) exceeds the number of sires. Using the conservative method of allele counting, Herridge (2016) found that *R. longicauda* females mated with a mean of 2.5 ± 0.15 SD (range 1-6) males. Similarly, our estimate of sire number, which was determined by counting the unique paternal alleles in each brood, averaged 3.6 ± 1.3 SD sires and ranged from 2-6. Based on this, the number of mates does not exceed the number of sires, suggesting that few or no males experience paternity failure in *R. longicauda*.

Our results are similar to paternity patterns found in two other species with valuable nuptial gifts: the ladybird beetle *A. bipunctata* (Haddrill et al. 2008) and orthopteran (katydid) *R. verticalis*, both with nutritious spermatophore meals (Simmons 2007). These insects showed high rates of multiple paternity (in 92.5%-100% of broods, respectively) and few males experienced paternity failure. In *R. verticalis*, analysis of stored sperm revealed that all males that inseminated a female sired offspring (Simmons 2007) and a similar pattern was evident for *A. bipunctata*, as the number of mates did not exceed the number of sires (Haddrill et al. 2008) (similar to our finding in dance flies).

By comparison, studies of insects that lack nuptial gifts and have no apparent sexual competition among females show much greater rates of paternity failure, as only 60% of a female’s mates sire offspring in the gryllid cricket *Telogryllus commodus*, 75% in *Telogryllus oceanicus* (Simmons and Beveridge 2010), and 51-66% in a third gryllid *L. cerasina* (Turnell and Shaw 2015). Together, these data support the hypothesis that paternity bias is reduced in species with valuable nuptial gifts. We suggest this is the best way of ensuring fitness (i.e., preventing complete paternity loss) when male mating frequency is constrained by long refractory periods and the fertilization success of investing males is limited by an inability to avoid sperm competition. Although shared paternity might reduce the number offspring for individual males, fitness costs are potentially reduced because frequently mated (fed-by-males) females may have higher fecundity or offspring quality (Browne 2021; Gwynne 1984b; reviewed in Arnqvist and Nilsson 2000; Vahed 1998).

Contrary to our prediction, a high degree of paternity bias was found in the nuptial gift-giving katydid, *P. griseoaptera* (Parker et al. 2017). However, as mentioned previously, this may also be due to the low percentage of a female’s lifetime production of offspring sampled. As the authors sampled less than 40% of a female’s lab-laid eggs, this may prevent successful sires from being detected (especially those hatched in future seasons), thus inflating the apparent rate of paternity failure.

In both our study species, there was significant paternity skew among sires, suggesting that offspring are not divided equally among successful males. This result is not unexpected, as it has been observed previously in gift giving insects *A. bipunctata* (Haddrill et al. 2008), *P. griseoaptera* (Parker et al. 2017), *E. ephippiger* (Hockham et al. 2004), and *R. verticalis* (Simmons 2007) as well as crickets that contribute little more than sperm when mating (*T. commodus, T. oceanicus*; Simmons and Beveridge 2010, and *L. cerasina*; Turnell and Shaw 2015). There are several possibilities that may explain paternity skew despite the prediction of shared paternity. First, random error during fertilization (e.g., slow or “sloppy” sperm mixing, sperm loss; Simmons 2001) is likely to cause small differences in paternity success among males that results in deviation from a perfect fair raffle (Herridge 2016). Because the degree of paternity skew was weak in both the ground weta and dance fly, the observed skew may not represent much more bias than would be expected at under a “noisy fair raffle” scenario (Herridge 2016). Alternatively, the observed skew could be caused by variation in number of sperm transferred (Simmons 2001; Parker et al. 2017). In *A. bipunctata*, Haddrill et al. (2008) used in-lab studies to determine that the observed skew among sires was due to differences in copulation duration. Differences in insemination duration may be especially pronounced in ground weta, since sperm transfer ends in ensiferan Orthoptera when the externally placed sperm ampulla (Brown and Gwynne 1997) is dislodged (male short-tailed ground weta may reduce incidences of dislodging by guarding their mates after copulation; Gwynne 2004).

Finally, we found no evidence of last male sperm precedence, a common paternity pattern in insects (Simmons 2001), in ground weta or dance flies. This further supports the hypothesis of reduced paternity bias and suggests that any observed skew was not due to mating order. The large, elastic sac of the spermathecae of ensiferan Orthoptera such as ground weta likely facilitates sperm mixing, even with frequent mating (Simmons 2001). In contrast, dance flies have a sclerotized, non-flexible spermathecae that would be expected to cause sperm displacement, with sperm from previous males being indirectly flushed from the storage organ (Simmons 2001). Indeed, Herridge 2016 found that stored sperm was dominated by a particular male in *R. longicauda* but this was not related to mating order or the resulting paternity shares. Importantly, one reason for the lack of last male precedence in dance flies observed in our study may be that our collection methods prevented this male from inseminating the female before uncoupling. However, at least one of the last male’s alleles were present in each clutch of offspring.

Additional constraints to our paternity analyses in dance flies are the incomplete genotypic data. Unlike in the ground weta, we were unable to genotype a female’s entire brood at all four loci, potentially reducing the accuracy of the parental reconstruction, as this limits the amount of information that can be used in the likelihood estimation (Jones and Wang 2010; Gerlach et al. 2012). This may explain the positive relationship between number of offspring analyzed and sires detected for dance flies. Further, we were unable to incorporate population allele frequencies into estimates of sire number (as in weta) because they did not include all alleles present in offspring (likely a consequence of high microsatellite polymorphism). Given these limitations, estimates of the most likely sire number in dance flies, including the offspring sired by the last male, may be subject to additional error. This may help to explain the high the upper limit on female mating rate (26 sires), however, this rate of polyandry is not unexpected given the biology of dance flies: Individually marked female *R. longicauda*, which may live several weeks (unpublished data), have been observed returning to (dawn and dusk) mating swarms, up to two times per day (R. Murray, personal communication).

Using microsatellite paternity analysis of wild-caught species where females rely on valuable nuptial gifts, we show that most males father offspring, resulting in low paternity bias (paternity sharing). We demonstrate this in species from two insect orders (Orthoptera and Diptera), using measures of lifetime female reproductive success in one to capture the final outcome of sperm competition. While sperm sharing reduces the number offspring for individual males, it reduces the chances of siring no offspring, which may be especially costly when males supply nuptial gifts. Thus, shared paternity, in which few males are excluded from siring offspring may make the evolution of large nuptial gifts more likely (Sakaluk 1986). These levels of paternity sharing may be expected in other species where females rely on costly mating gifts to produce offspring such as many katydids (Gwynne 1981; 1984b; 1985; 1988; Simmons and Bailey 1990) and other species of empidine dance flies (Diptera: Empididae; Downes 1970; Cumming 1994; Bussière et al. 2008; Wheeler et al. 2012; Murray et al. 2018).

## Acknowledgements

The New Zealand Department of Conservation kindly gave permissions to collect specimens. Genetic Marker Services developed of our microsatellite library and The Centre for Applied Genomics conducted genetic analyses. Thanks to: the Marc Johnson lab for use of equipment and other assistance, Lindsay Coome and Samreen Munim for their molecular work, Mary Morgan-Richards and Steve Trewick for advice, hospitality, and allowing us to collect, as well as Luc Bussière, Rosalind Murray, Marc Johnson, and Doug Currie) for comments on the research (funded by a NSERC Discovery Grant to DTG)

